# Warming-enhanced priority effects at population and community levels in aquatic bacteria

**DOI:** 10.1101/2020.01.27.921726

**Authors:** Máté Vass, Anna J. Székely, Eva S. Lindström, Omneya A. Osman, Silke Langenheder

## Abstract

The immigration history of communities can profoundly affect community composition. For instance, early-arriving species can have a lasting effect on community structure by reducing the immigration success of late-arriving ones through priority effects. Warming could possibly enhance priority effects by increasing growth rates of early-arriving bacteria. Here we implemented a full-factorial experiment with aquatic bacteria where both temperature and dispersal rate of a better-adapted community were manipulated to test their effects on the importance of priority effects, both on a community and a population level. Our results suggest that priority effects in aquatic bacteria might be primarily driven by niche preemption and strengthened by increasing temperature as warming increased the resistance of recipient communities against dispersal, and decreased the relative abundance of successfully established late-arriving bacteria. However, warming-enhanced priority effects were not always found and their strengths differed between recipient communities and dispersal rates. Nevertheless, our findings highlight the importance of context dependence of priority effects and the potential role of warming in mitigating the effects of invasion.

## Introduction

Variation in the composition of ecological communities can be the product of historical processes such as immigration, extinction and speciation (Fukami et al. 2007). The sequence and timing in which species or their propagules reach an ecological community (immigration history) can profoundly affect community structures and maintain diversity of communities via a process known as priority effects (Fukami 2015). Priority effects imply that early-arriving species gain advantage and become resistant to invasion of late-arriving ones, and therefore maintain high relative abundances over time (Lockwood et al. 1997). Priority effects are enforced by mechanisms that increase the ecological opportunity of the early-arriving species. For instance, factors such as time lag between arrivals, high growth and evolutionary rates of the early-arriving species (i.e., rapid local adaptation), are processes that reduce the establishment success of late-arriving species (De Meester et al. 2002, 2016). Successful local adaptation by early-arriving species initiates strong priority effects that can even reduce the establishment success of late-arriving species that are otherwise well-adapted to the local environment (Loeuille and Leibold 2008). On the other hand, priority effects can be absent or weak when the better-adapted late-arriving species generate species replacements. In the latter case, dispersal initiates species sorting processes, which has been shown to occur even at very low rates of dispersal (Declerck et al. 2013). In general, strong priority effects are expected if growth rates and/or the adaptive potential of local communities are high in relation to the time it takes for better-adapted species and/or dormant resident species to arrive or resuscitate and grow to become abundant community members (Vass and Langenheder 2017).

The most likely reason for priority effects is that early-arriving species induce rapid niche-modification or preempt resources so that late-arriving species will not be able to successfully establish in a local community (Tucker and Fukami 2014, Székely and Langenheder 2017). Hence, any possible environmental factor that increases growth rates of early-arriving species could possibly enhance priority effects (Chase 2010, Rudolf and Singh 2013). This includes the impact of ongoing climate change that leads to increased mean water surface temperatures (IPCC 2014). For instance, a recent study by Grainger et al. (2018) has shown that priority effects can be temperature-dependent in aphid species. In contrast, microorganisms are generally understudied in climate change-context studies (Cavicchioli et al. 2019) and it is currently unclear how the importance of priority effects for aquatic bacterial communities may be affected by warmer climate conditions.

Previous studies have demonstrated that organisms with high growth rates have the capability to facilitate strong priority effects (De Meester et al. 2002, Peay et al. 2012, Tucker and Fukami 2014), and therefore priority effects are expected to be more important in bacteria compared to other organism groups (De Meester et al. 2016). Several studies suggest priority effects occur in a variety of aquatic bacterial communities (Andersson et al. 2014, Svoboda et al. 2017, Rummens et al. 2018), but the effect of temperature on the process remains unexplored. Furthermore, we lack knowledge about the identity of bacteria that are key players in either causing priority effects or being hindered in their establishment by them. Even though two recent studies aimed to identify distinct roles of a bacterial groups in priority effects during community succession in biofilm (Brislawn et al. 2019) and in experimental freshwater bacterial communities (Rummens et al. 2018), it remains unclear whether the fates and roles of distinct bacteria in priority effects differ in response to warming.

Therefore, we performed a full-factorial experiment, where bacteria from three Swedish lakes were inoculated and grown in cell-free Baltic Sea medium at three temperature levels. These lake communities represented the early arrivals that were allowed to colonize the ‘foreign’ (Baltic Sea) medium to which they were not a priori adapted. After initial growth and establishment these communities became the ‘recipient communities’ that were exposed to invasion by Baltic Sea bacteria that were well-adapted to the incubation medium (i.e., Baltic Sea medium). These late-arriving communities were dispersed into the recipient communities at three different rates. We generated the recipient communities using three lake inocula that differed in their proximity to the Baltic Sea, as we expected that the distance of the origin of the early- and late-arriving communities might affect the strength of priority effects. At the end of the experiment, bacterial community composition was analyzed and the potential priority effects were investigated both at the community and population level.

We hypothesized that warming should result in stronger priority effects in bacterial communities, namely, that recipient communities grown at high temperature will be more resistant to dispersal of late-arriving species from the Baltic Sea. Moreover, we expected that recipient communities that are geographically closer to the Baltic Sea should harbor larger internal seed banks containing Baltic Sea bacteria and share the regional species pool with Baltic Sea to a greater extent than lakes further apart. Hence, we assumed that lakes geographically closer to the Baltic Sea would be more likely to maintain priority effects than more distant lakes due to their higher ratio of species adapted to the applied Baltic Sea medium.

## Material and methods

### Experimental design

In total, our experimental design resulted in 132 communities, three sets of recipient communities, exposed to three levels of dispersal and three temperatures, each with four replicates. For each temperature there was a dispersal source with four replicates and a control with four replicates consisting of cell-free medium (Fig. 1).

**Figure 1.**
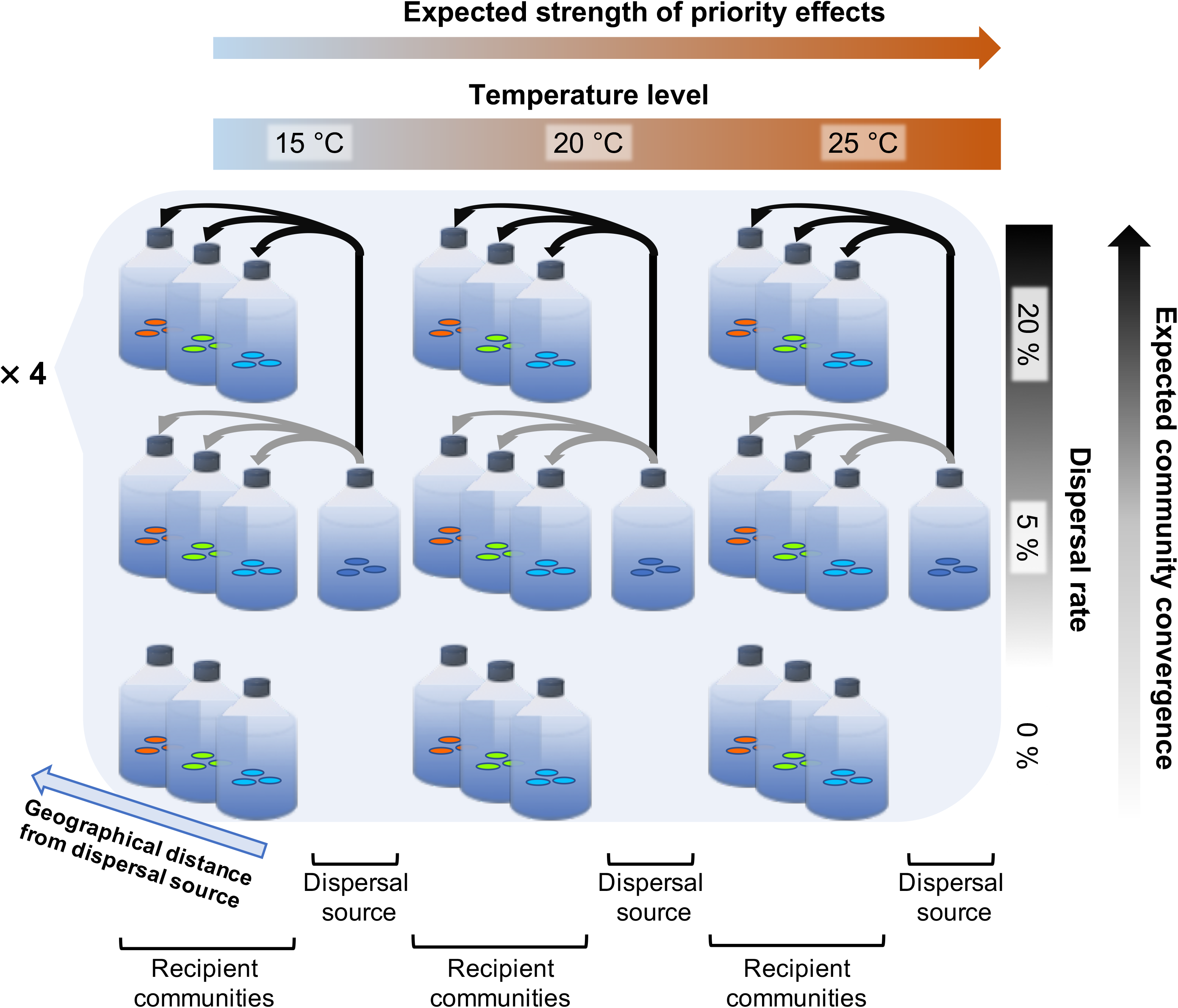
Experimental design of the study. The recipient communities were comprised of three different lake inocula (Erken, Lötsjön or Grytsjön, indicated by the different cell colors) inoculated separately into ‘foreign’ Baltic Sea incubation medium. The three lake inocula differed in their geographical distance from the Baltic Sea, with Grytsjön (in blue) being closest and Lötsjön (red) farthest away. The dispersal source constituted of the Baltic Sea community (dark blue cells) inoculated into cell-free incubation medium. Both the recipient (early-arriving lake bacteria) and the dispersal source (late-arriving Baltic Sea bacteria) communities were incubated at three different temperatures (15, 20 and 25 °C). Three different dispersal treatments (cell exchange) were applied by replacing 0, 5 and 20 % cells in the recipient communities with cells from the dispersal source. Black and grey arrows represent the direction of the dispersal treatments. The experiment was replicated four times. The recipient communities were always dispersed with the corresponding dispersal source replicate (n = 4) at the respective temperature level. Community convergence induced by dispersal was tested by measuring Bray-Curtis community dissimilarities between recipient and dispersal source communities as an indicator of the strength of priority effects: the less the communities converge toward the dispersal source, i.e., the more resistant recipient communities are against dispersal, the stronger are priority effects. The main hypothesis to be tested is that the strength of priority effects increases with temperature.

For the preparation of the Baltic Sea incubation medium used in this experiment, 120 L seawater was collected at the Swedish Baltic Sea coast on 19 June, 2018, at Barnens Ö (N 59°55’11.9”, E 18°54’52.2”). The water was filtered through a 20 μm net *in situ* to remove zooplankton and kept in dark at 4 °C overnight. Then, the medium was autoclaved (121 °C for 40 mins) and its pH was adjusted to its original level (pH = 8.18) by HCl addition. Afterwards, the medium was filtered through sterile 0.2 μm 47 mm membrane filters (Supor-200, Pall Corporation, Port Washington, NY, USA), and distributed into sterile 1,000 mL glass bottles, and autoclaved once more at 121 °C for 20 minutes in order to achieve a sterile cell-free incubation medium. Until inoculation, the bottles containing the sterile medium were kept in dark at 4 °C.

For the preparation of the inoculum communities, water samples were collected from three Swedish lakes (Lötsjön – N 59°51’44.0”, E 17°56’37.6”; Erken – N 59°50’09.2”, E 18°37’57.9”; and Grytsjön – N 59°52’21.1”, E 18°52’53.6”) and from the Baltic Sea (same location as above) on 4 July 2018 (Fig. S1). The distances of the three lakes from the Baltic Sea sampling location were 54.5 km (Lötsjön), 18.3 km (Erken) and 5.6 km (Grytsjön). The chemical characteristics of these lakes were slightly different; the concentrations of total carbon (TOC) and PO_4_^3−^ was higher, while NO_3_^−^ was lower in lakes located closer to the sea coast (Table S1). All samples were sequentially filtered to remove bacterial grazers, first through a 20 μm net *in situ* to remove zooplankton and then through GF/F filters (0.7 μm, Whatman, UK) prior inoculation to remove protozoans.

The dispersal source communities were established by inoculating 100 mL of GF/F filtered Baltic seawater into bottles containing 900 mL of cell-free Baltic Sea medium. The batch cultures were incubated at three different temperature levels (15, 20 and 25 °C) in the dark with four replicates at each incubation temperature. The established dispersal source communities were used in the dispersal treatments and represented the late-arriving species arriving at different rates (Fig. 1).

To create the recipient communities of early-arriving species 50 mL of GF/F filtered lake water inocula was added to bottles containing 450 mL cell-free Baltic Sea medium, and incubated at three different temperature levels (15, 20 and 25 °C) in dark with four replicates at each incubation temperature. The incubation of recipient cultures was started with one day delay so that cell abundance would most likely be lower compared to the dispersal sources, in order to avoid strong dilution of the medium during the dispersal process (see below).

### Dispersal

On day 7, after the successful establishment of recipient communities, measured as bacterial growth (Fig. S2), bacteria from the dispersal source communities were added to the recipient communities. The dispersal treatment consisted of one dispersal event at three different rates: no, low and high, wherein 0 %, 5 % and 20 % of the cells were exchanged with cells from the respective dispersal source (Fig. 1). For this, each replicate ‘A’ of the three recipient communities at the different incubation temperatures received cells from replicate ‘A’ of the dispersal source at the respective temperature level. Likewise, each replicate ‘B’ of the recipient communities received cells from replicate ‘B’ of the dispersal source and so on. For this, we measured the bacterial abundances (for details see ‘Sample analyses’ below) in all cultures and calculated the volume that needed to be replaced. To reach an equal final volume (564 mL) in all cultures the differences were compensated by adding additional cell-free incubation medium that was kept at the same conditions throughout the entire experiment. One ‘additional medium’ bottle (kept at 20 °C), broke during the experiment, hence, a mixture of the two other medium bottles (kept at 15 and 25 °C) were used after the dispersal treatments to reach equal volume in each incubation bottle. Both the cell exchange and the supply of additional medium were carried out under sterile conditions.

### Sample analyses

Throughout the experiment, bacterial abundance was monitored (Fig. S2) using a CytoFLEX flow cytometer (Beckman Coulter, Indianapolis, IN, USA) with 2.27 μM of SYTO 13 fluorescent nucleic acid stain (Invitrogen, Eugene, Oregon, USA).

To follow changes in environmental conditions in the cultures, samples for chemical analyses were collected three times: on day 1 after lake inocula were distributed into the medium, after the dispersal treatment (day 7), and on the last day of the experiment (day 22). Total phosphorus (TP), total nitrogen (TN) and total carbon (TOC) were measured spectrophotometrically (Perkin Elmer, Lambda 40, UV/VIS Spectrometer, Massachusetts, USA) and by catalytic thermal decomposition method (Shimadzu TNM-L, Kyoto, Japan), respectively according to standard procedures. Further, ion chromatography was used to measure the concentrations of NH_4_^+^, NO_3_^−^, PO_4_^3−^ as described previously (Attermeyer et al. 2019).

### Bacterial community composition

At the end of the experiment (day 22), the cultures (564 mL) were filtered by vacuum filtration onto 0.2 μm 47 mm membrane filters (Supor-200, Pall Corporation, Port Washington, NY, USA). DNA extraction from the membrane filters was performed using the DNeasy PowerSoil Kit (Qiagen, Venlo, Netherlands). The 16S rRNA gene amplicons were prepared using a two-step PCR protocol described in detail in the protocol deposited to the protocols.io repository (dx.doi.org/10.17504/protocols.io.6jmhck6). Amplicon paired-end sequencing was performed on Illumina MiSeq platform at the SciLifeLab SNP&SEQ Technology Platform hosted by Uppsala University, using Illumina MiSeq v3 sequencing chemistry. Raw sequences have been deposited to the European Nucleotide Archive with the accession number PRJEB34383.

Sequences were processed using DADA2 pipeline (Callahan et al. 2016) in R on the server of Uppsala Multidisciplinary Center for Advanced Computational Science (UPPMAX). First, forward and reverse sequences were trimmed to 280 and 220 bp long, respectively, after quality filtering (truncQ = 2) with maximum expected errors set to 2 and 5 for forward and reverse sequences, respectively. Secondly, sequences were dereplicated and sequence variants were inferred. Finally, chimeric sequences were removed and the final amplicon sequence variants (ASVs) were assigned against SILVA 132 core reference alignment (Quast et al. 2013).

### Data analyses

All statistical analyses and visualizations were conducted in R version 3.3.2 (R Core Team 2015). The ASV table was analyzed using the packages ‘phyloseq’ (McMurdie and Holmes 2013) and ‘vegan’ (Oksanen et al. 2016). Chloroplast ASVs and unassigned ASVs were discarded. Samples were rarified to an even depth of 6,366 reads per sample that eventually resulted in an ASV matrix with 5,598 ASVs in 120 samples. The taxonomic distribution of reads was visualized with Krona (http://sourceforge.net/projects/krona).

Principal component analysis was applied to assess if dispersal treatments induced any differences in nutrient concentrations during the experiment, including original, unfiltered lake and Baltic Sea water samples as references. Differences in bacterial abundance in dependence of temperature and the origin of the recipient community was tested using a two-way ANOVA and a subsequent Tukey’s HSD test.

Differences in community composition among samples were tested with permutational multivariate analysis of variance (PERMANOVA, permutations: 999) using the *adonis* function in ‘vegan’ package (Oksanen et al. 2016) and visualized using non-metric multidimensional scaling (NMDS), both based on the abundance-based Bray-Curtis dissimilarities. In the absence of priority effects, the well-adapted late-arriving bacteria from the Baltic Sea dispersal source should outcompete the originally maladapted early-arriving lake bacteria of the recipient community. Hence, the composition of the recipient communities would converge completely towards the dispersal source. On the opposite, we assumed the presence of priority effects when the recipient community maintained a significant dissimilarity compared to the dispersal source. To assess to what extent the recipient community shifted towards the dispersal source as a measure of the strength of priority effects, we first calculated the Bray-Curtis dissimilarity between each recipient community and its respective dispersal source. Our assumption was that priority effects are the stronger the higher the dissimilarity between recipient communities and the dispersal source. In case of complete priority effects at the level of the entire community, the recipient community should completely maintain its dissimilarity from the dispersal source. Thereafter, to be able to specifically address how dispersal influenced the strength of potential priority effects, we subtracted the Bray-Curtis dissimilarities between the recipient and dispersal source in the dispersal treatments with that of the respective 0 % dispersal treatment. This was done to correct for shifts in community composition that occurred in the recipient communities in the absence of dispersal.

Priority effects at the population level were investigated by determining the relative abundance of early-arriving lake ASVs of the recipient communities that persisted after exposure to dispersal, and late-arriving Baltic Sea ASVs of the dispersal source that established successfully in the recipient communities. For this, we first identified ASVs that were unique in either the recipient or the dispersal source communities, and fell in the above-mentioned categories by performing differential abundance analyses at each temperature level using the ‘DESeq2’ package (Love et al. 2014). First, we selected the non-common abundant ASVs (> 0.5 % relative abundance) in each recipient community and the dispersal source. Then, we determined separately for each recipient community whether the relative abundances (as a proxy for population size) of the abundant early-arriving ASVs changed after the effective dispersal treatments (i.e., 5 % and 20 % dispersal rate treatments) compared to their relative abundances in the no dispersal (0 %) communities. Here, we interpreted the lack of significant (adjusted *p* < 0.05) negative differences in relative abundances as a sign of priority effects and grouped the corresponding ASVs as ‘persistent early-arriving ASVs’. On the other hand, if their relative abundances were significantly lower (adjusted *p* < 0.05) in treatments receiving dispersal from the Baltic Sea, we categorized them as ‘forfeited early-arriving ASVs’. Second, for the late-arriving Baltic Sea ASVs in the dispersal source, we performed a conservative mixing analysis following Székely & Langenheder (2017). For this, we calculated the expected relative abundances of the abundant late-arriving ASVs’ (> 0.5 % in the dispersal source) in the effective dispersal treatments based on their relative abundances in the corresponding dispersal source, and the applied cell exchange rates (i.e., 5 or 20 %). For example, at 100 % efficacy, all dispersed late-arriving ASVs should have either 5 or 20 % of their relative abundances in the expected (recipient) communities. Thereafter we assessed the deviation of the measured abundances (i.e., in the recipient communities) from the calculated expected values. A non-significant deviation or a significantly (adjusted *p* < 0.05) higher relative abundance of a late-arriving ASV compared to the expected one provides a sign of successful establishment, while a significantly (adjusted *p* < 0.05) lower abundance indicates unsuccessful establishment of the late-arriving ASVs.

Finally, we used two-way ANOVAs with subsequent Tukey’s HSD test to assess if temperature and the origin of the recipient communities (inoculum origin) induced any differences on the relative abundance of persistent early-arriving ASVs and successfully established late-arriving ASVs.

## Results

After the initial inoculation of early-arriving bacteria in the Baltic sea medium all recipient communities showed typical growth patterns of dilution cultures and increased in abundance at least until day 7 (Fig. S2). The temperature increase (i.e., 20 and 25 °C) resulted in significantly higher abundances on day 7 compared to the 15 °C treatment (two-way ANOVA, F_Temperature_ = 76.09, *p* < 0.001; *post-hoc* Tukey’s HSD test: *p*_adjusted_ < 0.05, Table S2), further, there were significant differences based on the origin of the inoculum (F_Inoculum_ origin = 79.01, *p* < 0.001) and the interaction between inoculum origin and temperature was also significant (F_Temperature × Inoculum origin_ = 7.86, *p* < 0.001) on day 7. Bacterial abundances remained stable throughout the experiment in all treatments. Despite some initial variation, the chemical conditions of the cultures inoculated with different recipient communities did not experience any pronounced shift or showed clustering patterns in response to the dispersal treatments (Fig. S3A) or to the different incubation temperatures (Fig. S3B).

The NMDS of the bacterial communities (Fig. 2) and PERMANOVA shows that without dispersal (i.e., 0 % dispersal rate) all three recipient communities (Lötsjön, Erken and Grytsjön) were compositionally different (Fig. 2 orange dots; PERMANOVA: F_Inoculum origin_ = 9.96, R^2^ = 0.35, *p* = 0.001), and were affected by the temperature manipulation (F_Temperature_ = 3.02, R^2^ = 0.11, *p* = 0.001). The recipient communities exposed to dispersal (i.e., 5 and 20 % dispersal rate, brown and black dots) became more similar to the dispersal source (Fig 2; blue dots). However, PERMANOVA results showed that the recipient communities exposed to dispersal were significantly dissimilar from the dispersal source in all cases (Table S3), thus, complete convergence to the dispersal source (i.e., complete absence of priority effects) did not occur in any of the communities. We found a similar pattern when assessing the degree of dissimilarity of recipient communities from the dispersal source. More specifically, recipient communities receiving either 5 % or 20 % dispersal shifted towards the dispersal source, without a complete convergence (Bray-Curtis dissimilarity ≠ 0) (Fig. 3).

**Figure 2.**
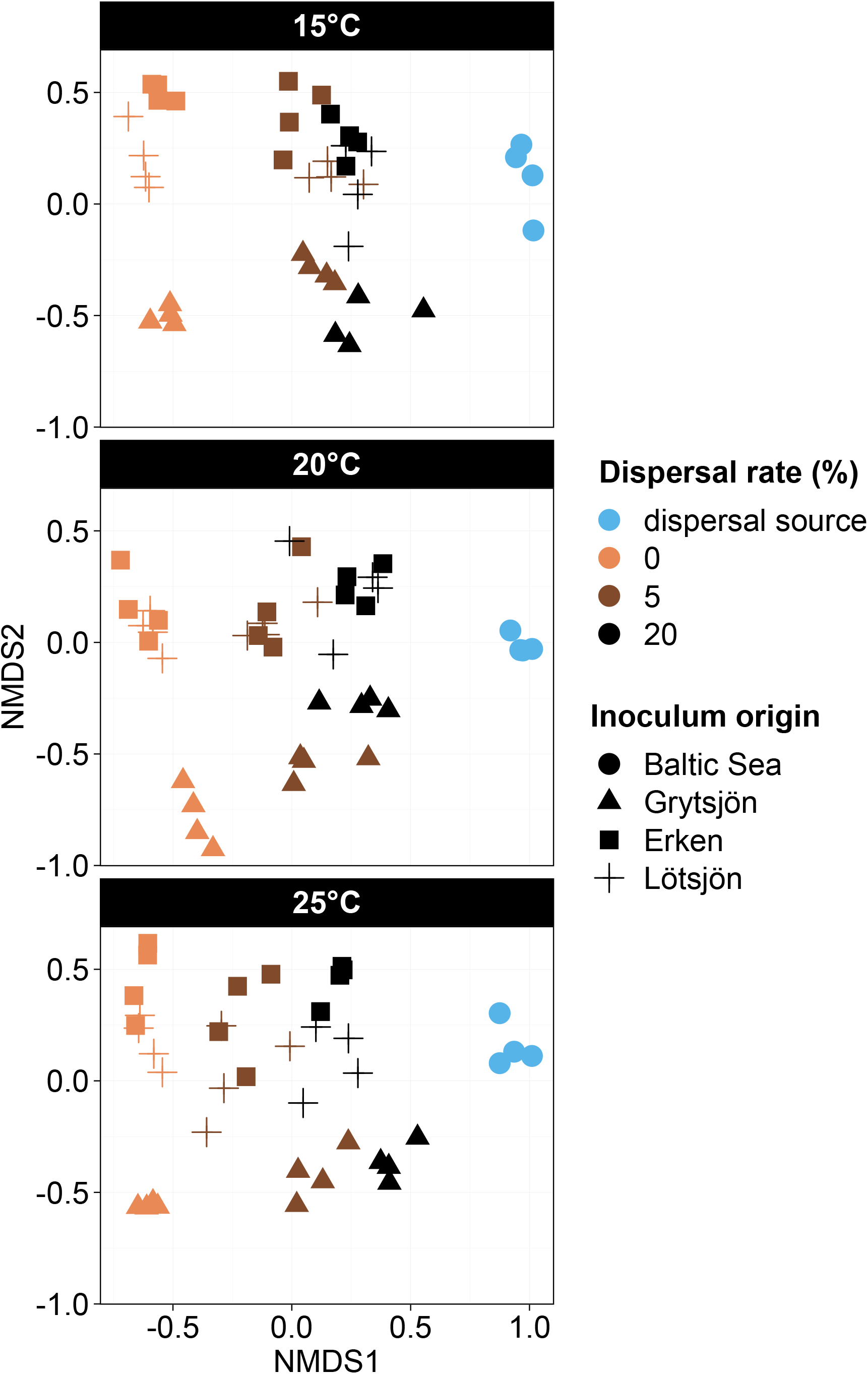
Non-metric multidimensional scaling (NMDS) plots derived from abundance-based Bray-Curtis dissimilarities of bacterial community composition at the three temperature levels by the end of the experiment. Note that cultures with Baltic Sea inoculum were used as the dispersal source, while cultures with lake inocula (Grytsjön, Erken and Lötsjön) were used as recipient communities. All cultures were grown in Baltic Sea medium. Symbols are shaped and colored by inoculum origin and dispersal treatment, respectively. Goodness of fit (stress value): 0.105.

**Figure 3.**
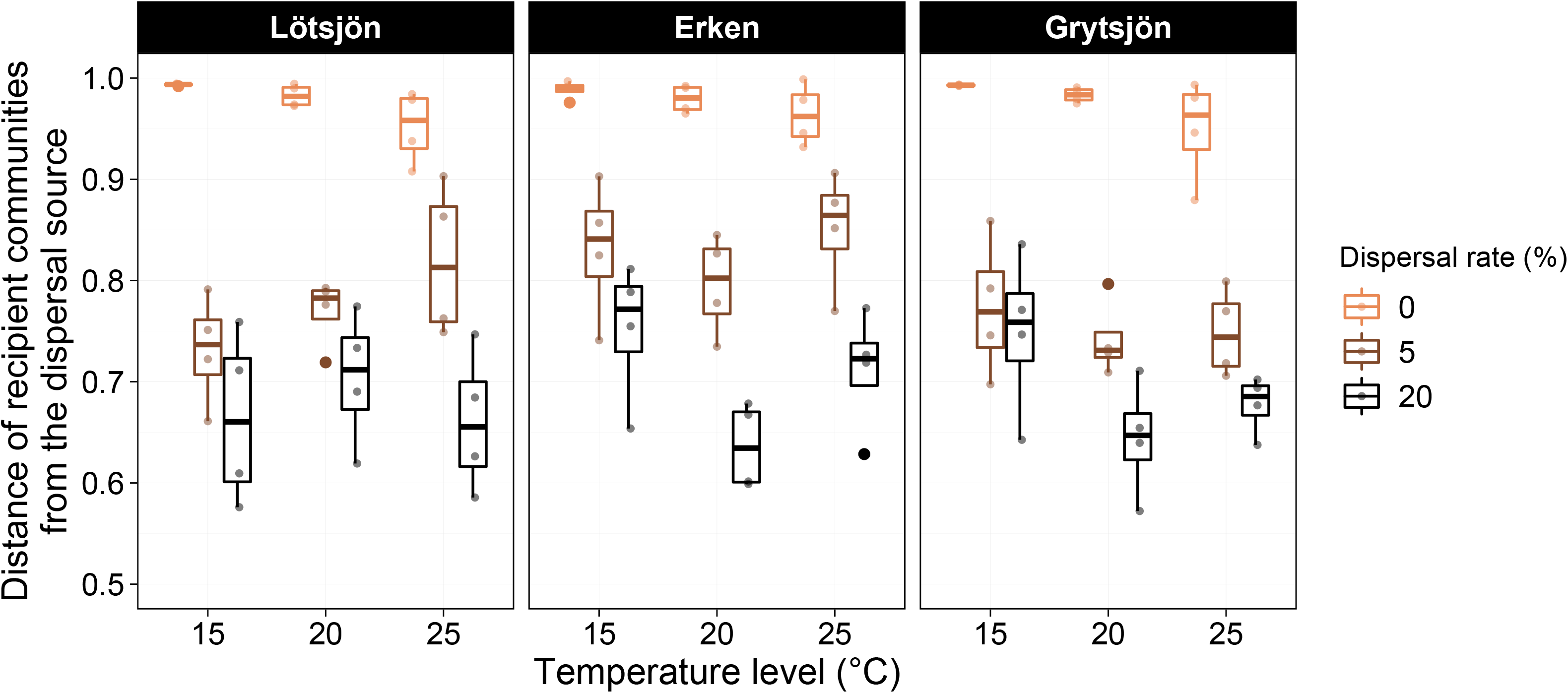
Distance of recipient communities from the dispersal source (based on Bray-Curtis dissimilarity) at different temperature levels. Recipient communities were exposed to either 0, 5 or 20 % dispersal (cell exchange) from the dispersal source.

Recipient communities with the highest dispersal (i.e. 20 %) shifted the most towards the dispersal source (Fig. 3). When Bray-Curtis dissimilarities were calculated in relation to the 0 % dispersal treatment to assess the degree of community shift induced by dispersal (as a proxy for the strength of the priority effects), the shift was greater in the 20 % dispersal compared to 5 % dispersal rate treatment (Fig. 4), i.e. priority effects were stronger at 5 % dispersal compared to 20 %. Both temperature and the inoculum origin of the recipient community had a significant effect on this relationship at 5 % dispersal (two-way ANOVA, F_Temperature_= 5.97, *p* = 0.006, F_Inoculum origin_= 5.76, *p* = 0.007), but not at 20 % dispersal (two-way ANOVA, F_Temperature_ = 1.99, *p* = 0.153, F_Inoculum origin_ = 0.33, *p* = 0.724) (Fig. 4). This indicates that the strength of priority effects was affected by temperature and recipient community origin at 5 % dispersal but not at 20 %. More specifically, stronger priority effects occurred (i.e., less community shift induced by dispersal) at higher temperature. There were no significant interactions between temperature and inoculum origin in any of the dispersal rates tested.

**Figure 4.**
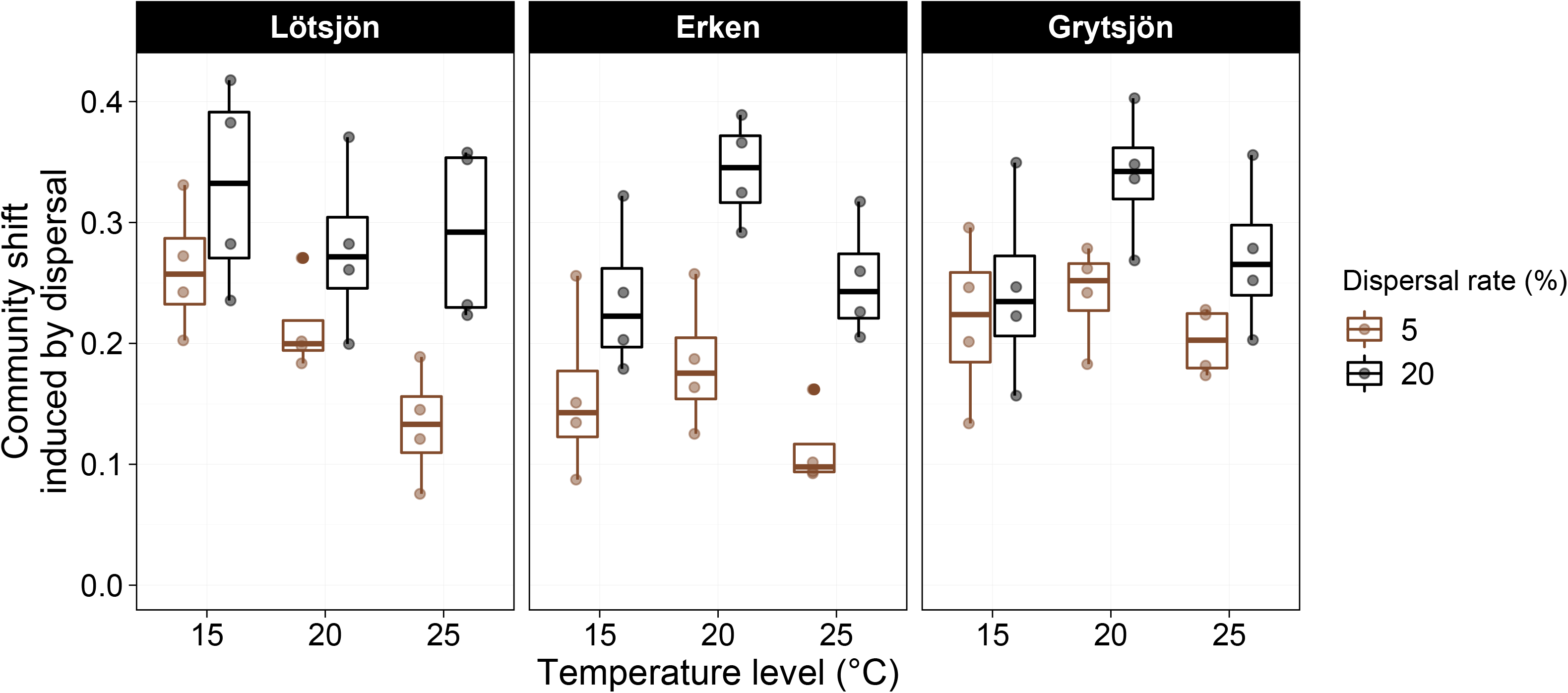
Dispersal-induced shifts in community composition in relation to temperature. Differences in Bray-Curtis dissimilarities between recipient communities and the dispersal source in the 5 and 20 % dispersal treatments in relation to the 0 % dispersal treatment.

We further examined changes in the dynamics of early- and late-arriving ASVs in response to temperature changes. On a broad taxonomical level, we found that the abundant (> 0.5 % relative abundance) bacterial ASVs in the early-arriving communities belonged to the class Alphaproteobacteria, Gammaproteobacteria and Bacteroidia (Fig. S4). The most abundant genera (top three) were *Brevundimonas*, *Pseudomonas*, *Allorhizobium-Neorhizobium-Pararhizobium-Rhizobium* (thereafter *A-N-P-R*) in the Lötsjön and Erken recipient communities and *Limnobacter*, *Algoriphagus* and *A-N-P-R* in the Grytsjön recipient communities (Fig. S4). The abundant (> 0.5% relative abundance) members of the dispersal source communities (i.e., late-arriving bacteria) were ASVs belonging to Alphaproteobacteria (mainly *Loktanella*, *A-N-P-R* and *Roseibacterium*), Gammaproteobacteria (mainly *Hydrogenophaga*, *Pseudomonas*, *Rheinheimera*) and Bacteroidia (mainly *Algoriphagus*) (Fig. S4, Baltic Sea).

Differential abundance analyses revealed numerous ASVs of the abundant genera (> 0.5 %) that could be classified as either ‘persistent’ or ‘forfeited’ early-arriving ASVs or ‘successful’ or ‘unsuccessful’ late-arriving ASVs (see Methods). We identified several persistent early-arriving ASVs (belonging to the genera of *A-N-P-R*, *Brevundimonas*, *Flavobacterium*, *Limnobacter*, *Novosphingobium*, *Perlucidibaca*, *Pseudomonas*, *Rheinheimera* and *Sphingorhabdus*) that occurred in all three recipient communities. In addition, ASVs of *Fluviicola* and *Hydrogenophaga* were also persistent (i.e., did not show significant (*p*_adjusted_ < 0.05) changes in relative abundance in the dispersal treatments) in Erken and Grytsjön recipient communities, but not in Lötsjön (Fig. 5, S5). Several ASVs within the above-mentioned genera were forfeited by the dispersal of late-arriving Baltic Sea bacteria, and in Lötsjön *Sphingobium* represented the only genus containing merely forfeited ASVs found. Interestingly, most of the forfeited ASVs occurred in Lötsjön (n = 27) and Grytsjön (n = 51) recipient communities, while only two forfeited ASVs (*Sediminibacterium* grouped as ‘other_Bacteroidia’) occurred in Erken recipient communities. There were also inconsistences because ASVs from the same genera (e.g. *Brevundimonas, Pseudomonas* and *Flavobacterium*) could be categorized both as forfeited and persistent early-arriving bacteria.

**Figure 5.**
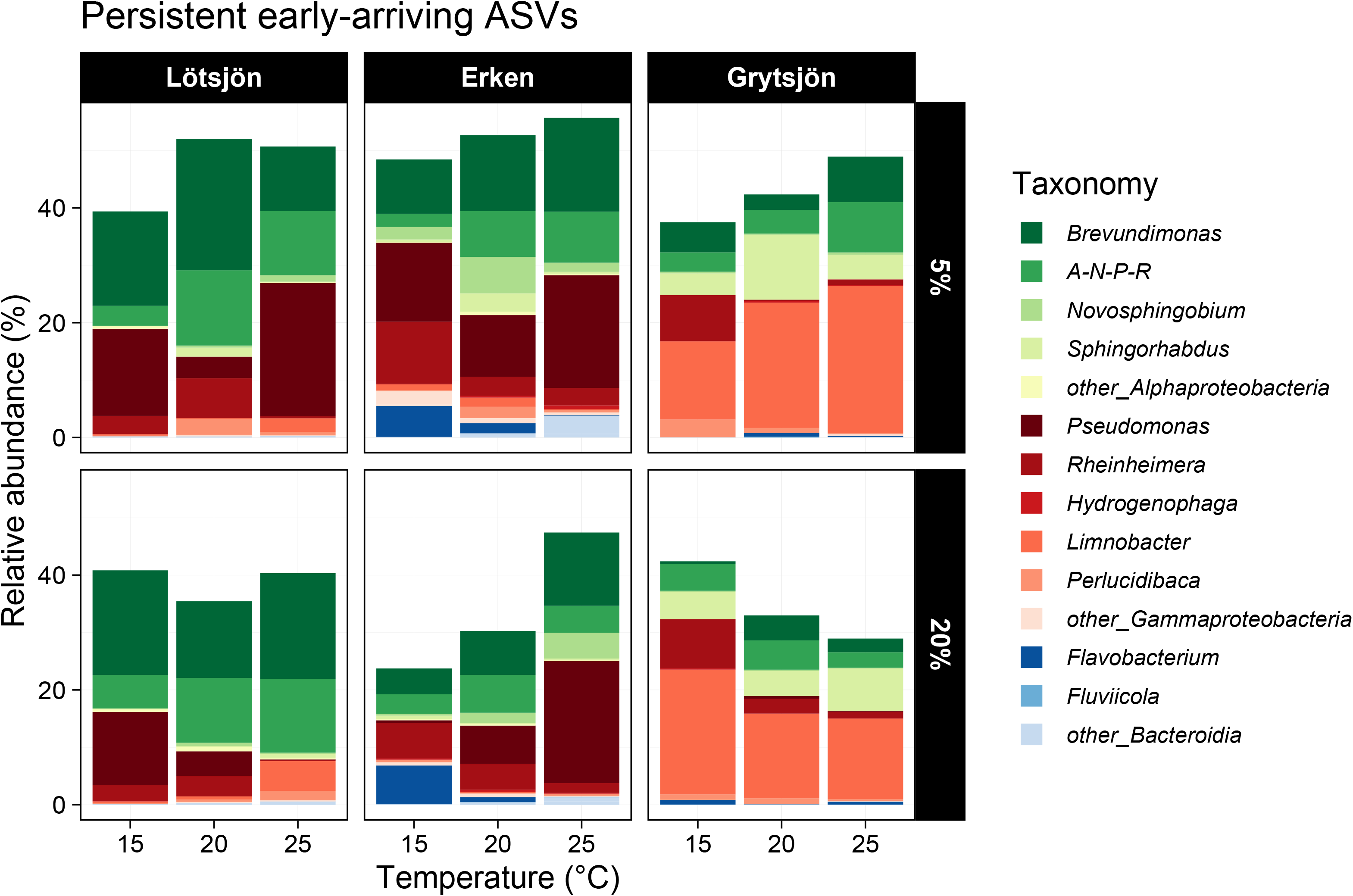
Changes in the relative abundances of abundant persistent early-arriving bacteria (ASVs > 0.5 % relative abundance) in the different dispersal (5 % or 20 %) and temperature treatments (15, 20 and 25 °C). ASVs are grouped by bacterial genus and were identified by differential abundance analysis (see Methods for the assessment procedure and Figure S6 for further results). *A-N-P-R* refers to the genus *Allorhizobium-Neorhizobium-Pararhizobium-Rhizobium*.

Interestingly, changes in the composition of persistent early-arriving ASVs were found as the temperature level increased. For example, there was a general trend showing that *Flavobacterium* and *Rheinheimera* were more persistent at lower temperature (15 °C and 20 °C) than at the highest temperature level (25 °C). In contrast, *A-N-P-R*, *Brevundimonas* (especially in the case of Erken), *Fluviicola* and non-abundant groups (i.e. *Sediminibacterium, Chryseolinea and* unknown Bacteroidia grouped as ‘other_Bacteroidia’; *Comamonas* and *Herbaspirillum* grouped as ‘other_Gammaproteobacteria’; *Ferrovibrio* grouped as ‘other_Alphaproteobacteria’in Fig. 5) tended to be more abundant and persistent at higher temperatures. Temperature and inoculum origin had, however, no effect on the total relative abundance of the persistent early-arriving ASVs in the recipient communities with different dispersal treatments (two-way ANOVA at 5 % dispersal: F_temperature_ = 2.38, *p* = 0.109, F_inoculum origin_ = 1.95, *p* = 0.159; 20 % dispersal: F_temperature_ = 0.74, *p* = 0.488, F_inoculum origin_ = 0.59, *p* = 0.562, no significant interactions in either case) (Fig. 5).

Among late-arriving Baltic Sea bacteria, mainly ASVs of *Algoriphagus*, *Loktanella*, *Roseibacterium*, *Hydrogenophaga* were the ones that showed successful establishment, thus, maintained or significantly increased (adjusted *p* < 0.05) their relative abundances after being dispersed into recipient communities (Fig. 6, Fig. S6). The composition of successfully established late-arriving bacteria was similar regardless of the recipient communities into which they were dispersed. Their total relative abundances differed across temperature levels in both the 5 % and 20 % dispersal treatments and differed between recipient communities with different inoculum origin (two-way ANOVA at 5 % dispersal: F_temperature_ = 7.98, *p* = 0.002, F_inoculum origin_ = 3.83, *p* = 0.033; at 20 % dispersal: F_temperature_ = 7.34, *p* = 0.002, F_inoculum origin_ = 4.79, *p* = 0.015, no significant interactions in either case) (Fig. 6). Significant differences (based on post-hoc Tukey tests at *p* < 0.05) were found between 15 vs. 25 °C and 20 vs. 25 °C, respectively, and between Erken and Grytsjön at 5 % dispersal and Lötsjön and Grytsjön at 20 % dispersal. Among the populations most impacted by warming were *Loktanella*, *Hydrogenophaga* and *Pseudomonas* which showed decrease in relative abundance at higher temperature levels (20 and 25 °C) (Fig. 6, Fig. S6).

**Figure 6.**
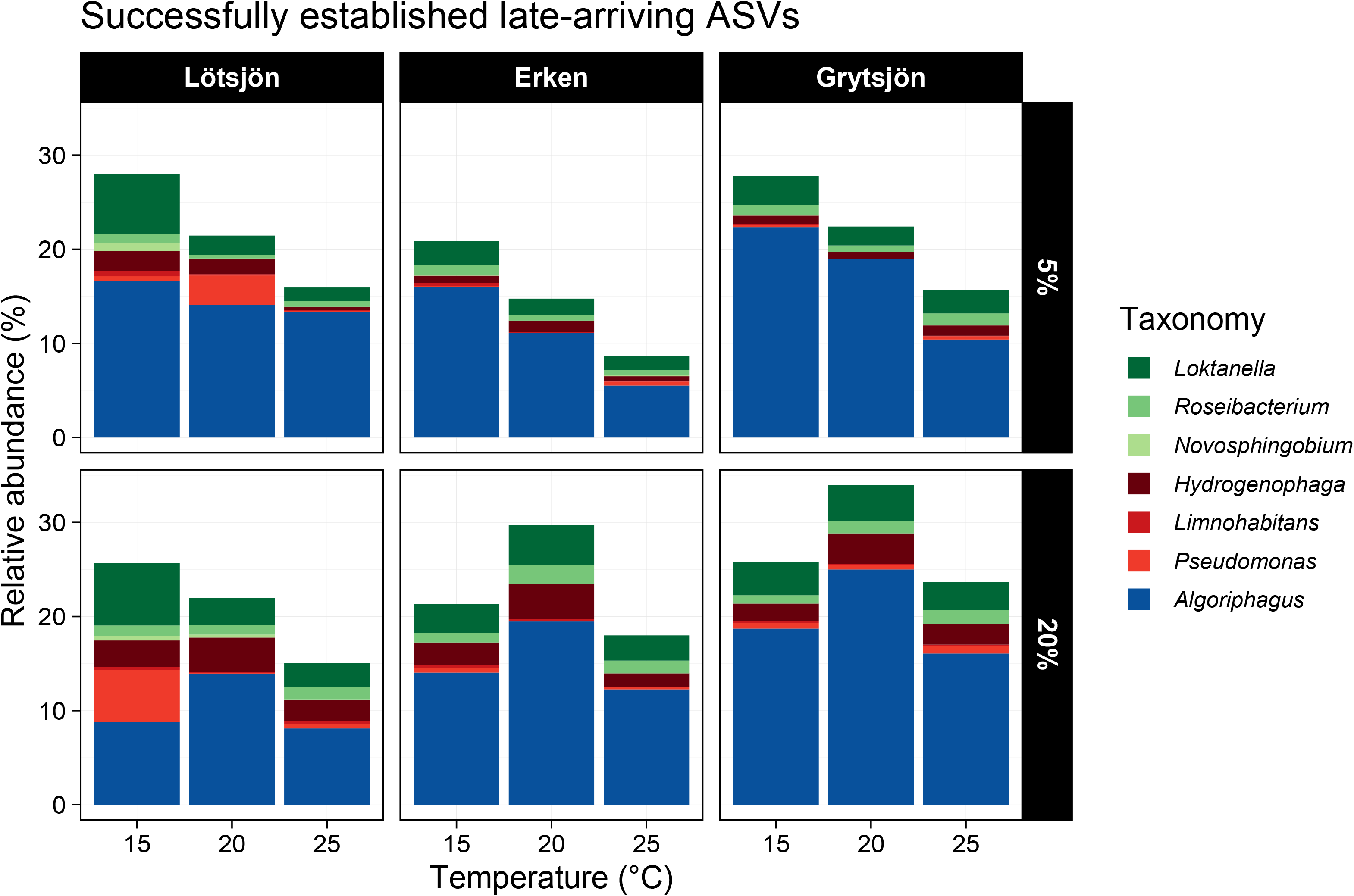
Changes in the relative abundances of successfully established late-arriving species (ASVs, > 0.5 % relative abundance) in the different dispersal (5 % or 20 %) and temperature treatments (15, 20 and 25 °C). ASVs are grouped by bacterial genus and were identified by differential abundance analysis (see Methods for the assessment procedure and Figure S7 for further results).

## Discussion

We found evidence of priority effects in aquatic bacteria in agreement with previous studies that have used similar approaches to ours (Svoboda et al. 2017, Rummens et al. 2018). More specifically, we showed that warming has the potential to promote priority effects, but that it depends (i) on the dispersal rate of late-arriving better-adapted communities into recipient communities and (ii) on the composition of the recipient community. We also found that dispersal of the late-arriving bacteria induced species replacement, i.e. decreased priority effects, because all recipient communities converged towards the dispersal source communities to some extent.

Previous studies have primarily focused on priority effects related to individual species or strains (Fukami et al. 2007). However, in our study we tried to mimic natural dispersal events of bacterial communities which are often complex and involve mixing or coalescence of entire communities (Rillig et al. 2015). Hence, to our knowledge, this study represents the first experimental evidence that temperature-dependency of priority effects can occur in complex pelagic bacterial communities wherein different bacterial groups are involved in different ways. The warming effect could be seen at the population level since less successful establishment of late-arriving ASVs were found in the recipient communities in response to increasing temperature. Specifically, in the case of the recipient communities, the total relative abundance of successfully established ASVs generally decreased with increasing temperature, whereas the relative abundance of persistent early arriving ASVs tended to show the opposite trends, even though this was not significant in any case.

One possible explanation for the lower establishment success and stronger persistence of resident species at higher temperatures is that the resistance of recipient communities to invasion (dispersal) by late-arriving bacteria increased as a result of temperature-stimulated high growth rates of the early-arriving bacteria (see Fig. S2 and Table S2). Similarly, Grainger et al. (2018) demonstrated that increased temperature increased growth rates of aphid species, thus, allowing them to more rapidly change and deplete resources which altogether increased the competitive exclusions of competitor species that arrived late. In our experiment, such effects appeared to be generally stronger at 5 compared to 20 % dispersal rates. This highlights that dispersal rates are an important mediator of the strength of priority effects in natural communities, and that this strength in general is likely to be higher if dispersal rates of late arrivals are relatively low (Loeuille and Leibold 2008).

We identified several persistent early-arriving ASVs that taxonomically differed between the three sets of recipient communities. Hence, distinct sequence variants (ASVs) of early-arriving bacteria played a role in maintaining priority effects. However, we also found that there can be inconsistencies at the genus level in the response to dispersal of different sequence variants as ASVs belonging to the very same genus (e.g., *Pseudomonas, Flavobacterium* and *Rheinheimera*) can be categorized both as persistent and forfeited ASVs. Since similar results have been obtained in a number of studies (Fukami et al. 2007, Tucker and Fukami 2014, Rummens et al. 2018, Brislawn et al. 2019) of different complexity, our results suggest that species’ responses to invasion at the population level are difficult to predict. Nevertheless, our finding corroborates results of other recent studies (Needham et al. 2017, García-García et al. 2019) that emphasize the importance to evaluate population level dynamics at the deepest taxonomical resolution possible. Moreover, the composition of persistent early-arriving ASVs differed between the different temperature treatments, suggesting that, as temperature conditions change, the identity of bacterial populations that maintain priority effects changes as well. On the other hand, in the case of the dispersal source community, we found consistency in the identity of the successfully established late-arriving ASVs as, irrespective of the identity of the recipient community or the temperature treatment, they typically belonged to the same genera. In summary, our findings suggest that different species can be involved in the development of priority effects of aquatic bacterial communities under different circumstances.

The lake inocula included for the preparation of the recipient communities in this study differed in their geographical distance to the dispersal source (the late-arriving bacteria from the Baltic Sea). Therefore, we presumed that recipient communities closer to the Baltic Sea might have been exposed to dispersal from the Baltic Sea in their recent history to a greater extent than those farther from the Baltic. This could have resulted in a larger shared species pool, including larger numbers of bacteria of Baltic Sea origin in local lake seed banks (Comte et al. 2014). We therefore presumed that the potentially higher numbers of species that are adapted to environmental conditions in the Baltic Sea in recipient communities closer to the Baltic would lead to stronger priority effects. Our results do, however, not support this idea because the dissimilarities between the recipient and dispersal source communities were similarly high in all cases (see Fig. 3). Hence, the results do not support our hypothesis of stronger priority effects in lakes closer to the Baltic Sea. Further, it suggests that the shared species pools with the Baltic Sea, including local seed banks of Baltic Sea taxa, were equally low irrespective of the distance of the lake to the Baltic Sea. There were nevertheless differences among lakes regarding the pattern of how temperature affected dispersal-induced shifts in community composition at the community level and the total relative abundance of persistent early-arriving ASVs and successfully established late-arriving ASVs. These differences might be the consequence of the differences of chemical characteristic of the three lakes (Table S1), the compositional differences of the initial bacterial communities (Figure S4), or the result of intrinsic differences in traits (e.g. temperature optima) of ASVs that contribute to priority effects in the different lakes that we cannot disentangle in our study.

Priority effects can be due to two distinct mechanisms: niche-modification and niche preemption (Fukami 2015), but providing insights into the mechanisms underlying priority effects is difficult. In our experiment niche modification-driven priority effects of the different lake inocula should have influenced the identities of the successfully established late-arriving bacteria, which was, however, not the case. On the contrary, niche preemption might have had an influence in communities grown at increased temperatures (20 and 25 °C) as communities had attained higher abundances at the time when the dispersal source was added, while the establishment success of late-arriving ASVs decreased with increasing temperatures (see Fig. 6). This indicates that the availability of resources was probably reduced to such an extent in 20 and 25 °C treatments that this limited the abundances of late-arriving bacteria.

In addition to warming and dispersal there might be other factors that can influence the importance of priority effects in natural bacterial communities. For instance, it remains unclear what would happen in the presence of predation (e.g., bacterial grazers) or multi-level trophic interactions. A few previous studies on zooplankton communities suggested that predation can be an important factor and can either reduce priority effects (Louette and De Meester 2007, Berga et al. 2015), or, in contrast, indirectly promote them (Ryberg et al. 2012). However, we lack a comprehensive knowledge on how predation could affect priority effects in particular in microbial communities. Another aspect that need to be considered are temperature fluctuations that can promote the immigration success of dispersed species and maintain multiple species coexistence, thus, reducing historical contingency (Litchman 2010, Tucker and Fukami 2014, Toju et al. 2018).

Organisms across multiple kingdoms might be negatively affected by global warming, altering their ecosystem functions (Altermatt et al. 2008, Hall et al. 2008, Rudolf and Singh 2013, Dong et al. 2018). Temperature has been shown to stimulate microbial invasions (e.g., spread of vibrios; Vezzulli *et al.* 2012) and influence the biogeographical patterns of microbes (Amalfitano et al. 2014). However, priority effects could play an important role by dampening the establishment success of invasive bacteria (Thomsen et al. 2011). Our experimental study also shows the potential invasion mitigating effect of warming as with increasing temperature priority effect of early-arriving aquatic bacteria by niche preemption increasingly hindered the establishment success of late-arriving bacteria from an external dispersal source. However, our findings also highlight that the overall strength of warming-enhanced priority effects is context dependent and differs depending on the composition of early-arriving communities as well as dispersal rates of the late-arriving ones.

## Acknowledgements

We thank Vasiliki Papazachariou for her help and assistance during field sampling and the implementation of the experiment. We are also grateful for Christoffer Bergvall for his help in the chemical analyses of samples. This research was supported by grants from the Swedish Research Council to S.L., the Malméns Foundation to M.V. and the Swedish Research Council Formas to A.J.S..

## Data accessibility

The sequencing data supporting the results are archived in the public repository European Nucleotide Archive with accession numbers PRJEB34383. Additional data (e.g. ASV tables, sample and chemical data) are available on the openly available repository of Uppsala University (DiVA; http://urn.kb.se/resolve?urn=urn:nbn:se:uu:diva-409821) under the following ID: diva2:1427510.

## Author contributions

S.L., M.V., A.J.S. and E.S.L. designed the study. M.V. performed the experiment. O.A.O. processed the samples. M.V. performed data processing, analyses and drafted the manuscript. All others contributed substantially to writing and revisions.

