## Supplementary Material for "Warming-enhanced priority effects at population and community levels in aquatic bacteria"

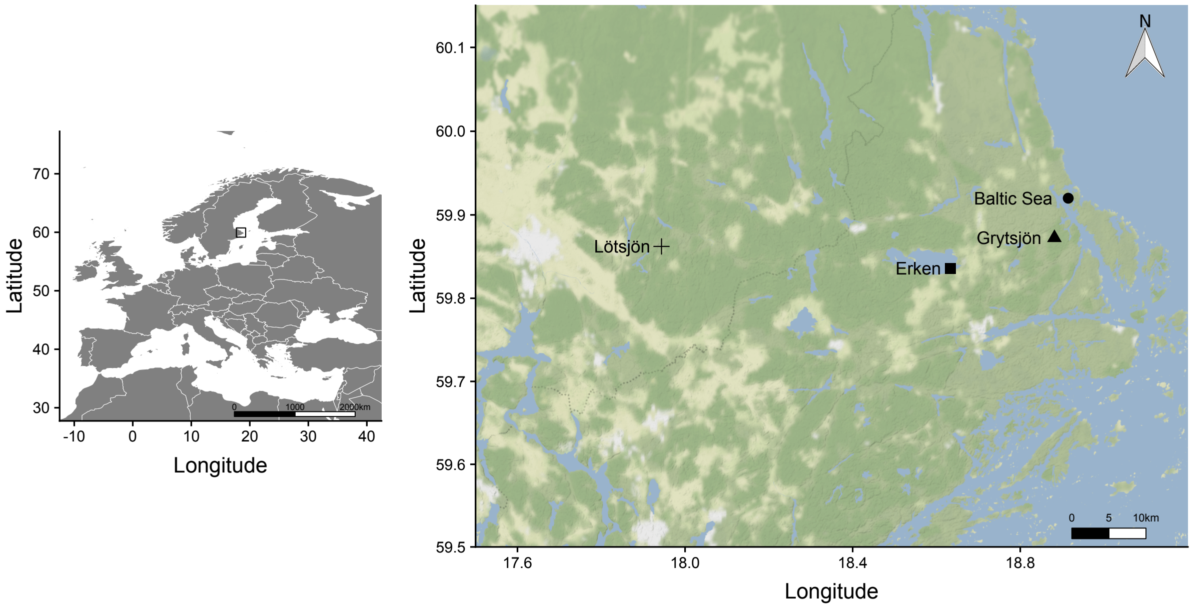


**Figure S1.** Map of sampling locations. Samples for the recipient communities were collected from lakes Lötsjön, Erken and Grytsjön, and the dispersal source inoculum as well as water for the preparation of the Baltic Sea medium was collected from the Baltic Sea at the coast of Barnens Ö, Sweden. Map tiles by Stamen Design, under a Creative Commons Attribution license (CC BY 3.0). Data by OpenStreetMap, under the Open Data Commons Open Database License (ODbL).

**Table S1.** Average chemical characteristics of the sampling sites. TP: total phosphorous, TN: total nitrogen, TOC: total organic carbon.

| | **Lötsjön** | **Erken** | **Grytsjön** | **Baltic Sea** |
|---|---|---|---|---|
| TN [mg/L] | 0.596 | 0.772 | 0.656 | 0.159 |
| TP [mg/L] | 16.185 | 15.992 | 19.268 | 16.185 |
| TOC [mg/L] | 10.240 | 11.495 | 14.365 | 3.871 |
| $NH_4^+$ [µg/L] | <5 | <5 | <5 | 10.326 |
| $NO_3^-$ [µg/L] | 27.438 | 19.403 | 15.495 | 5.419 |
| $PO_4^{3-}$ [µg/L] | 6.861 | 9.245 | 12.653 | 3.388 |

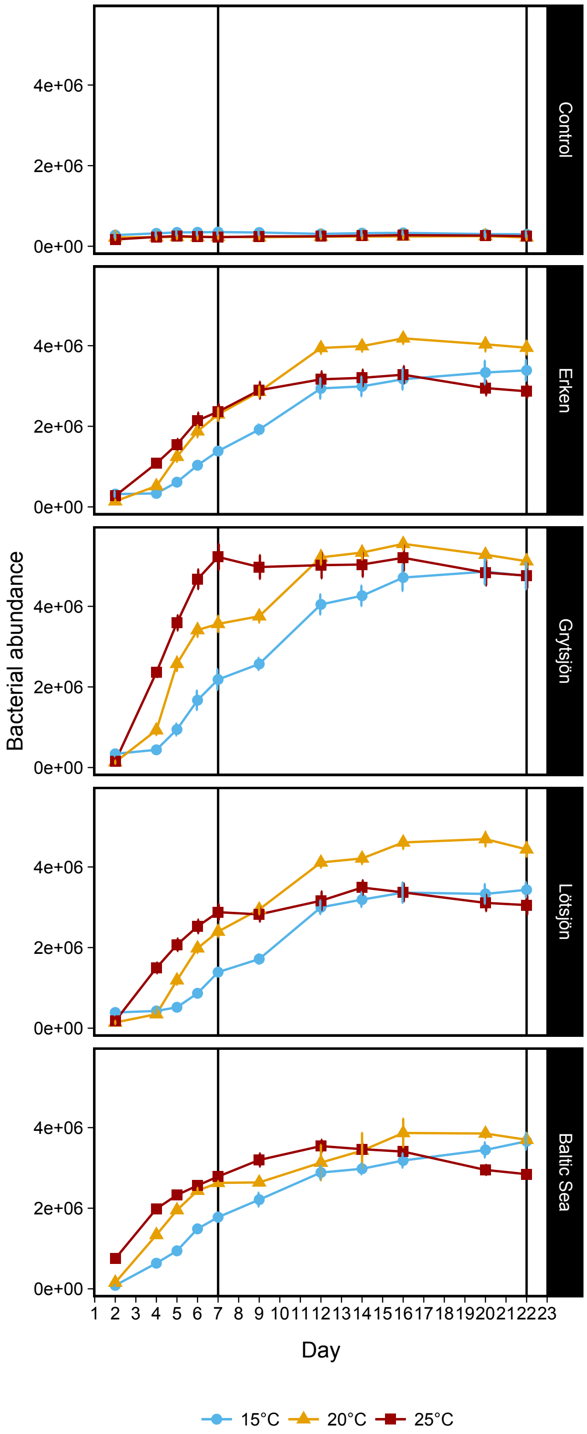


**Figure S2.** Growth curves of bacterial communities in the cultures incubated at three temperature levels (15, 20 and 25 °C). The first vertical line (day 7) represents the time point at which bacteria from the dispersal source were added to the recipient communities, while the second one, on day 22, shows the end of the experiment. N = 4 for each type of culture and treatment at each date. Controls consisted of non-inoculated incubation medium.

**Table S2.** Results of a *post-hoc* Tukey's HSD test assessing differences in bacterial abundances on day 7 between cultures grown at different temperature levels (15, 20 or 25 °C).

| | **adjusted *p*** |
|---|---|
| **Lötsjön** | |
| 15 – 20 | **7.79E-03** |
| 15 – 25 | **2.40E-06** |
| 20 – 25 | 8.34E-01 |
| **Erken** | |
| 15 – 20 | **2.79E-02** |
| 15 – 25 | **1.18E-02** |
| 20 – 25 | 1.00E+00 |
| **Grytsjön** | |
| 15 – 20 | **1.81E-05** |
| 15 – 25 | **0.00E+00** |
| 20 – 25 | **1.00E-07** |

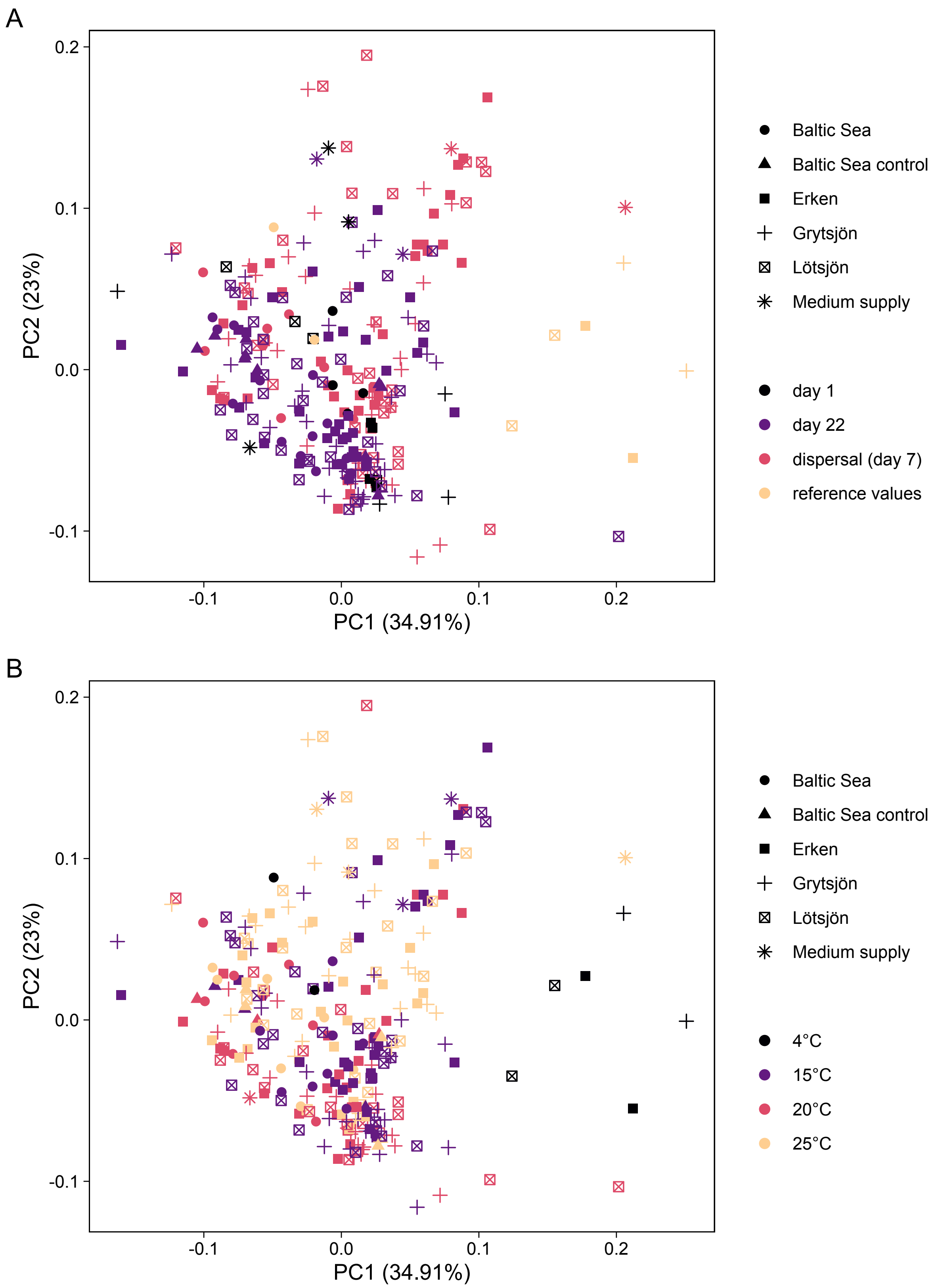


**Figure S3.** Principal component analysis showing differences in the chemical conditions ($NH_4^+$, $NO_3^-$, $PO_4^3$, TOC, TN and TP) between (A) the different cultures and samples, and (B) different temperature levels. Note that the reference values, i.e., the natural unprocessed lake (Erken, Lötsjön and Grytsjön) and seawater (Baltic Sea) samples were kept at 4 °C before the medium and inocula preparations.

**Table S3.** Results of the PERMANOVA models testing the dissimilarity between recipient community (early-arriving bacteria) and dispersal source (late-arriving bacteria) at different temperature levels and dispersal rates. Bold values indicate significance at $p < 0.05$.

| Lake compared to the dispersal source | F value | $R^2$ | *p* | Temperature (°C) | Dispersal rate (%) |
|---|---|---|---|---|---|
| Lötsjön | 5.392 | 0.473 | **0.033** | 15 | 0 |
| | 2.470 | 0.292 | **0.029** | 15 | 5 |
| | 2.293 | 0.276 | **0.028** | 15 | 20 |
| | 12.970 | 0.684 | **0.024** | 20 | 0 |
| | 6.273 | 0.511 | **0.024** | 20 | 5 |
| | 3.396 | 0.361 | **0.043** | 20 | 20 |
| | 7.615 | 0.559 | **0.031** | 25 | 0 |
| | 3.298 | 0.355 | **0.029** | 25 | 5 |
| | 3.430 | 0.364 | **0.028** | 25 | 20 |
| Erken | 7.579 | 0.558 | **0.031** | 15 | 0 |
| | 2.868 | 0.323 | **0.033** | 15 | 5 |
| | 3.137 | 0.343 | **0.028** | 15 | 20 |
| | 7.963 | 0.570 | **0.026** | 20 | 0 |
| | 6.082 | 0.503 | **0.020** | 20 | 5 |
| | 3.857 | 0.391 | **0.030** | 20 | 20 |
| | 9.597 | 0.615 | **0.022** | 25 | 0 |
| | 5.313 | 0.469 | **0.033** | 25 | 5 |
| | 3.904 | 0.394 | **0.041** | 25 | 20 |
| Grytsjön | 7.174 | 0.545 | **0.042** | 15 | 0 |
| | 3.531 | 0.370 | **0.032** | 15 | 5 |
| | 3.235 | 0.350 | **0.028** | 15 | 20 |
| | 12.710 | 0.679 | **0.034** | 20 | 0 |
| | 6.573 | 0.523 | **0.021** | 20 | 5 |
| | 5.073 | 0.458 | **0.028** | 20 | 20 |
| | 8.550 | 0.588 | **0.022** | 25 | 0 |
| | 6.881 | 0.534 | **0.037** | 25 | 5 |
| | 4.150 | 0.409 | **0.023** | 25 | 20 |

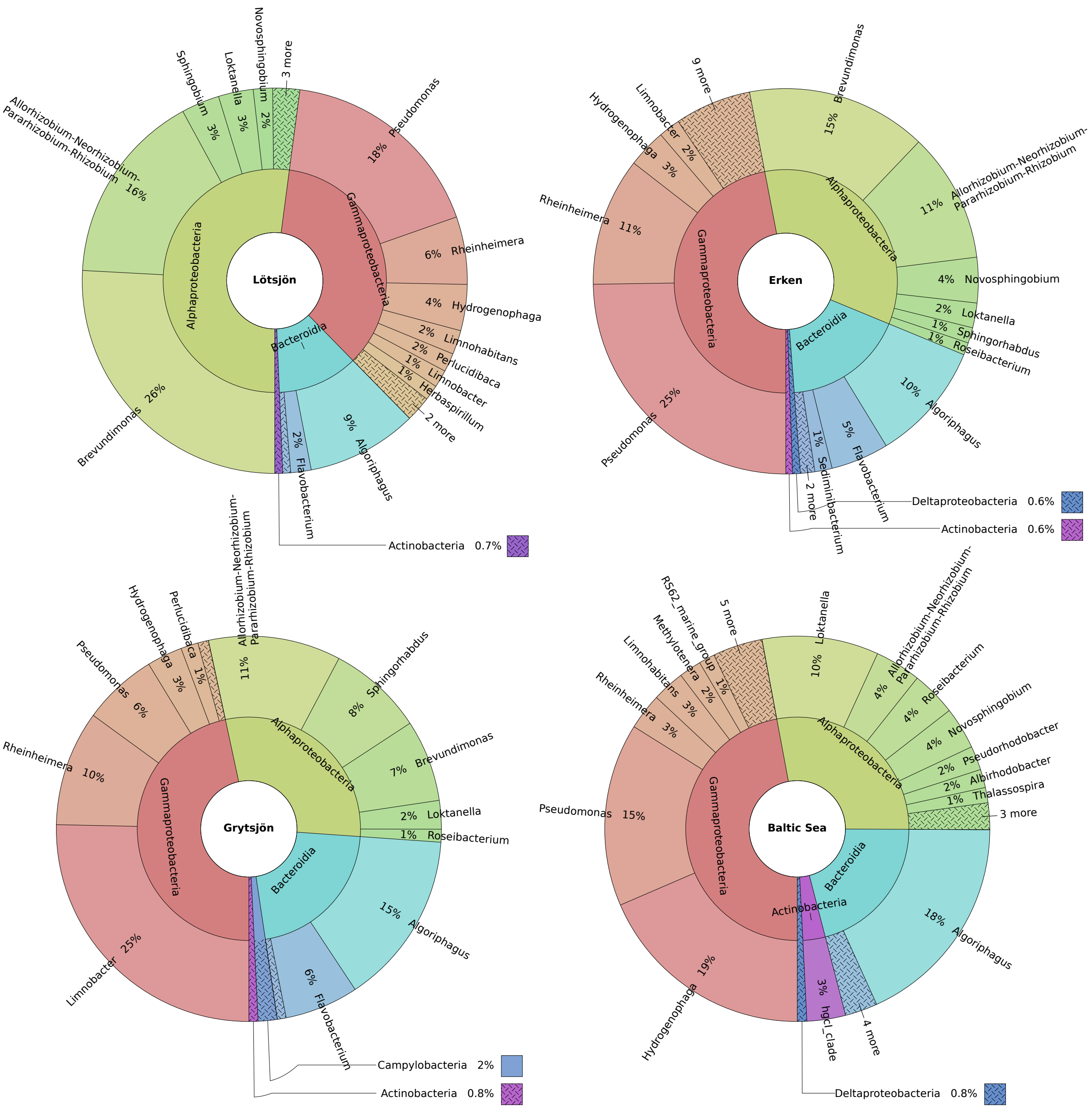


**Figure S4.** Taxonomic distributions of the most common bacterial genera (> 0.5 % relative abundance of ASVs) found in the cultures with different initial inoculum.

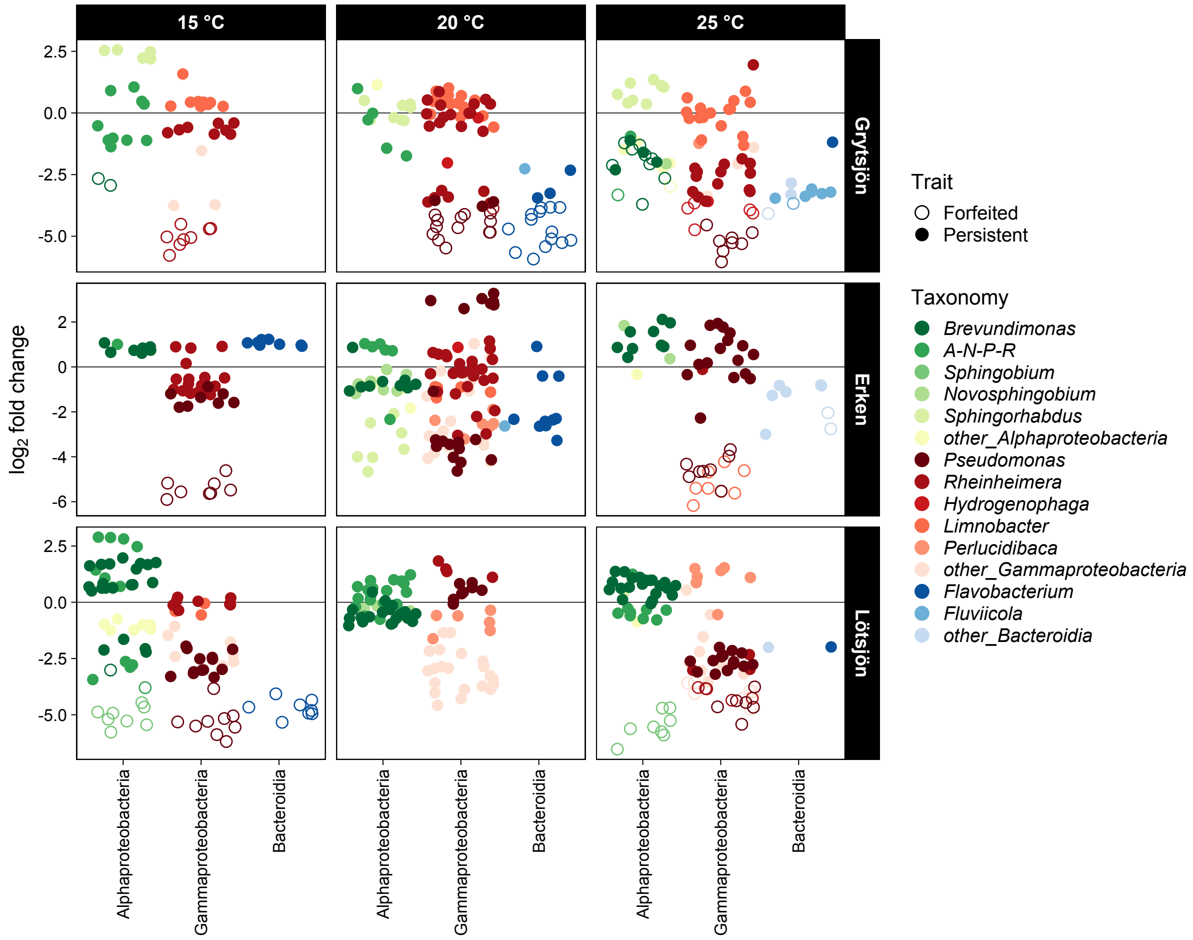


**Figure S5.** Differential abundance analyses of early-arriving bacteria (16S amplicon sequence variants, ASVs) of recipient communities comparing communities with 0 % and 5 or 20 % dispersal rates, respectively. Non-significant and positive (adjusted $p < 0.05$) $\log_2$ fold change values (filled dots) indicate similar or higher abundance when late-arriving bacteria were introduced and , hence, ASVs are defined as persistent, while negative values indicate ASVs with significantly lower abundance (empty dots), and, hence, are defined as forfeited. *A-N-P-R* refers to the genus *Allorhizobium-Neorhizobium-Pararhizobium-Rhizobium*.

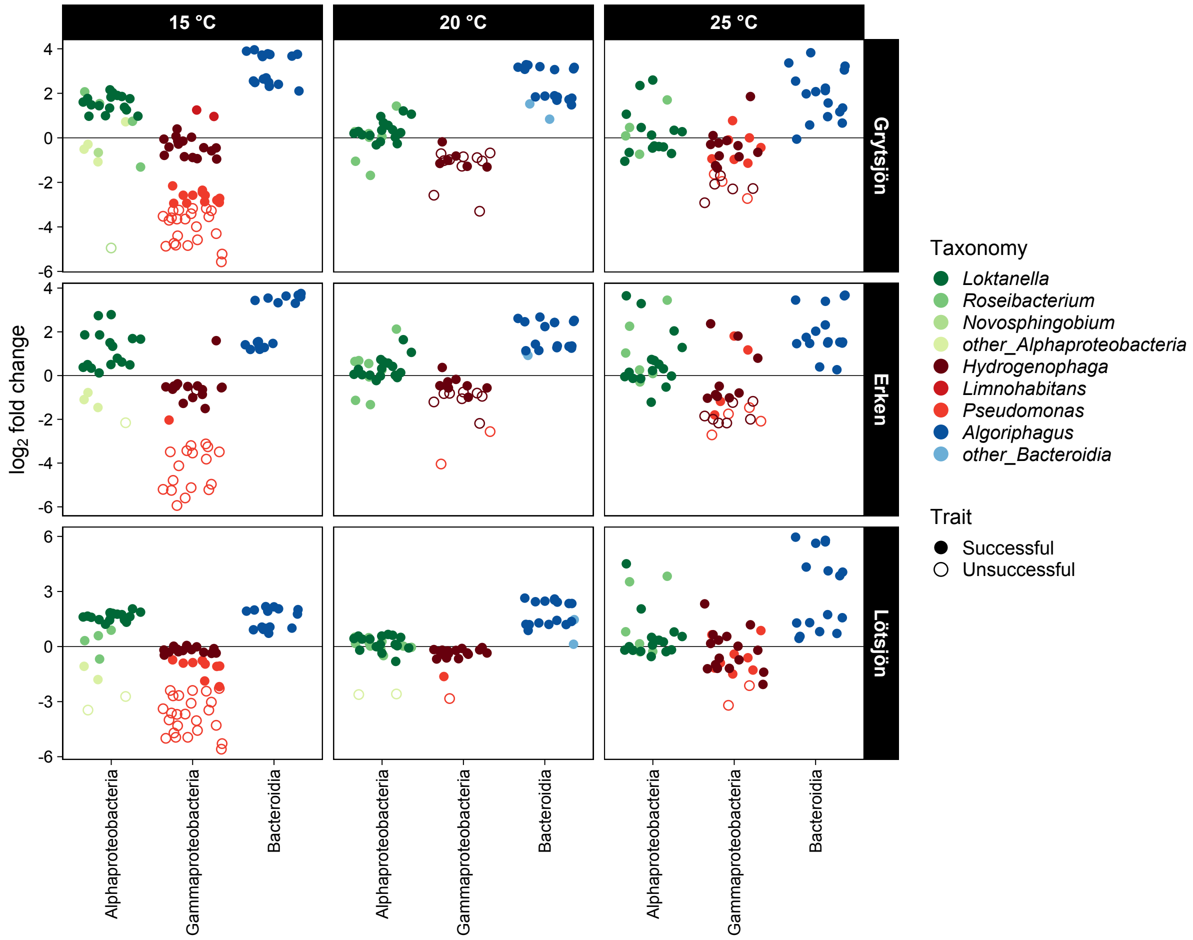


**Figure S6.** Differential abundance analyses of late-arriving bacteria (16S amplicon sequence variants, ASVs) comparing the expected and actual abundance of late-arriving ASVs in recipient communities with 5 and 20 % dispersal rates. Non-significant and positive (adjusted $p < 0.05$) $\log_2$ fold change values (filled dots) indicate similar or higher abundance, hence, successfully establishment of ASVs, while negative values indicate significantly lower abundance (empty dots) and, hence, unsuccessful establishment of ASVs after being dispersed.
